# Widespread noncoded amino acids in human proteome

**DOI:** 10.1101/292474

**Authors:** Jing-Hua Yang, Xinjun Chen, Jing Gong, Xin Lv, Han Zhao, Cuiling Li, Baibing Bi, Fengqin Wang, Shengnan Sun, Xingyuan Wang, Haobo Zhang, Tao Huang, Kazem Azadzoi, Feng Shi, Xianglong Kong, Minglei Shu, Yinglong Wang, Wan Huang, Y. Eugene Chin, Zhinan Chen, Zi-Jiang Chen

**Affiliations:** State Key Laboratory of Microbial Technology, Cancer Research Center, Schools of Basic Medical Sciences, Life Sciences and Management, Shandong University, Jinan, China; Departments of Surgery and Urology, VA Boston Healthcare System, Boston University School of Medicine, Boston MA, USA; Center for Reproductive Medicine, the Key laboratory for Reproductive Endocrinology of Ministry of Education, Shandong Provincial Hospital, Shandong University, Jinan, China; Shandong Institute for Food and Drug Control, Jinan, China; Shandong Computer Science Center (National Supercomputer Center in Jinan), Shandong Provincial Key Laboratory of Computer Networks, Jinan, China; National Molecular Medicine Science and Technology Center, the State Key Laboratory of Cancer Biology, The Fourth Military Medical University, Xi’an, China; The Key Laboratory of Stem Cell Biology, Institutes of Health Sciences, Shanghai Institutes for Biological Sciences, Chinese Academy of Sciences, Shanghai, China; Center for Reproductive Medicine, Shanghai Key Laboratory for Assisted Reproduction and Reproductive Genetics, Ren Ji Hospital, Shanghai Jiao Tong University School of Medicine, Shanghai, China

**Keywords:** Proteomics, post-translational modification, mass spectrometry, amino acid substitution, amino acid polymorphism, sperm proteome, oligoasthenospermia, Gaussian clustering

## Abstract

Proteins are usually deciphered by translation of the coding genome; however, their amino acid residues are seldom determined directly across the proteome. Herein, we describe a systematic workflow for identifying all possible protein residues that differ from the coding genome, termed noncoded amino acids (ncAAs). By measuring the mass differences between the coding amino acids and the actual protein residues in human spermatozoa, over a million nonzero delta masses were detected, fallen into 424 high-quality Gaussian clusters and 571 high-confidence ncAAs spanning 29,053 protein sites. Most ncAAs are novel with unresolved side-chains and discriminative between healthy individuals and patients with oligoasthenospermia. For validation, 40 out of 98 ncAAs that matched with amino acid substitutions were confirmed by exon sequencing. This workflow revealed the widespread existence of previously unreported ncAAs in the sperm proteome, which represents a new dimension on the understanding of amino acid polymorphisms at the proteomic level.

**Highlights:**

- 571 ncAAs spanning 108,000 protein sites were identified in human sperm proteome.
- Most ncAAs are novel with unresolved sidechains and found at unreported protein sites.
- Exon sequencing confirmed 40 of 98 ncAAs that matched with amino acid substitutions.
- Many ncAAs are linked with disease and have potential for diagnosis and targeting.

**eTOC Blurb:** We describe a systematic identification of all possible protein residues that were not encoded by their genomic sequences. A total of 571 high-confidence most novel noncoded amino acids were identified in human sperm proteome, corresponding to over 108,000 ncAA-containing protein sites. For validation, 40 out of 98 ncAAs that matched to amino acid substitutions were confirmed by exon sequencing. These ncAAs are discriminative between individuals and expand our understanding of amino acid polymorphisms in human proteomes and diseases.

## INTRODUCTION

Proteins are polyamides that are encoded by their corresponding genes and built from 20 different types of amino acid. Protein residues are often subjected to post-translational modifications (PTMs), which are involved in regulation of protein folding, stability and activity, and ultimately their function (Beltrao et al., 2012; Mathias et al., 2015). However, protein sequences are usually deciphered by translation of the coding genomic sequences and the actual protein residues are seldom determined directly across the proteome (Chick et al., 2015). Thus, the actual structures of residues in proteins are assumed to be more complicated than thought. This is partially because protein sequencing technologies are usually dependent on proteins translated from their encoding genes (Mann and Wilm, 1994; Perkins et al., 1999). The leading Shotgun proteomics technology, for instance, begins with enzymatic digestion of proteins, followed by mass determination of the digested peptides (precursors or MS), and collision-induced fragmentation of the peptides (products or MS/MS) using liquid chromatography-tandem mass spectrometry (LC-MS/MS). Peptide sequences are determined by matching the obtained MS and MS/MS spectra with the theoretical mases of translated or coding protein peptides using sophisticated search algorithms such as MASCOT (Smith and Frank, 2016), SEQUEST (Lundgren et al., 2009), and more recently, Byonic (Bern et al., 2012). In matching algorithms, only the standard 20 amino acids and known modifications are considered, and undefined or noncoded protein residues that do not match these standards are usually neglected. Surprisingly, nearly half of high-quality MS/MS spectra do not match peptides in the available protein databases (Chick et al., 2015). Even when considering known modifications, many spectra still do not match the coding proteins due to undefined noncoded protein residues.

To resolve unmatched peptide spectra, the existence of one or more undefined noncoded residues in peptides is often assumed. In early approaches, partial peptide sequences were derived from unmatched spectra and used as tags to search theoretically translated protein databases, thereby identifying unexpected PTMs and amino acid substitutions (Mann and Wilm, 1994; Tabb et al., 2003). In iterative approaches, partially matched or de novo-generated mass tags are used to test mass iterations for multiple unexpected modifications (Bern et al., 2007; Kim et al., 2009). Alternatively, unrestrictive search algorithms can identify noncoded residues without knowing whether or not they exist in nature (Tsur et al., 2005). The mass-tolerant approach was originally used to detect known modifications by allowing for mass differences between a precursor and its fragment (Creasy and Cottrell, 2002). This approach was recently improved by allowing a wide mass tolerance for precursors to match peptide sequences containing a wide range of delta masses or undefined modifications (Chick et al., 2015). Although the high rate of false positives is still a major problem of these approaches, along with variable sensitivity and lengthy experiments, it is clear that unmatched peptide spectra are largely due to yet undefined noncoded protein residues. Since undefined noncoded amino acids indicate novel amino acid side-chains and structures, they are assumed to be important for protein function under both physiological and pathological conditions

Sperm are highly differentiated cells in which many genes are transcriptionally and translationally silent. This study explored noncoded protein residues in human spermatozoa, with an emphasis on comparing healthy individuals and patients with oligoasthenospermia. Taking advantage of the high accuracy and fast data acquisition of mass spectrometry, a modified Shotgun proteomics method was used to measure all possible mass differences or delta masses between coding amino acids and actual protein residues in sperm proteins from 48 people. Over a million nonzero delta masses were obtained out of ~7 million peptides or 10,591 non-redundant proteins detected with high confidence. To account for the false positive rate, nonzero delta masses were clustered by a multi-variable model and further fitted by Gaussian regression to archive statistically clustered delta masses for calculation of high-confidence ncAAs and ncAA-containing protein sites. Our study revealed a large number of unreported ncAAs with undefined side-chain structures in human spermatozoa that are polymorphous in healthy populations and discriminative in patients with oligoasthenospermia.

## RESULTS

### Widespread and highly clustered delta masses in human sperm proteome

In this study, delta masses refer to mass differences between genetically encoded amino acids and the actual protein residues at the corresponding peptide position. The existence of “nonzero” delta masses indicate the presence of potential noncoded protein residues or noncoded amino acids, termed ncAAs. Positive and negative delta masses indicate mass gain and loss, respectively. A modified Shotgun proteomics approached was used to determine delta masses for spermatozoa proteins from 24 healthy individuals and 24 patients with oligoasthenospermia. The total proteins were separated into 5-10 fractions based on molecular weights by SDS-PAGE and in-gel trypsin digestion. Peptides (precursors, MS) and their breakdown fragments (products, MS/MS) were repeatedly determined 3-8 times by liquid chromatography-tandem mass spectrometry (LC-MS/MS) to maximise the number of high-quality peptide spectrum data. By allowing noncoded protein residues in peptides, the acquired MS and MS/MS data were matched against human protein databases using the multi-blind spectral alignment algorithm MODa (Na et al., 2012), wide-tolerance SEQUEST (Chick et al., 2015), and Byonic (Bern et al., 2007). All three algorithms detected a large number of peptides with nonzero delta mass (**Fig. 1a**). For Byonic, 1,197,754 nonzero delta masses were detected (**Supplemental Table 1**) out of 7,860,173 high-confidence peptides, indicating that approximately 15% of the peptides potentially included at least one ncAA in their sequence. All peptides were grouped into 10,591 non-redundant proteins (at least two peptides were required to identify a protein) with an average of 26.75% sequence coverage (**Fig. 1b**), representing the highest number and coverage of proteins identified from human spermatozoa to date.

**Fig. 1.**
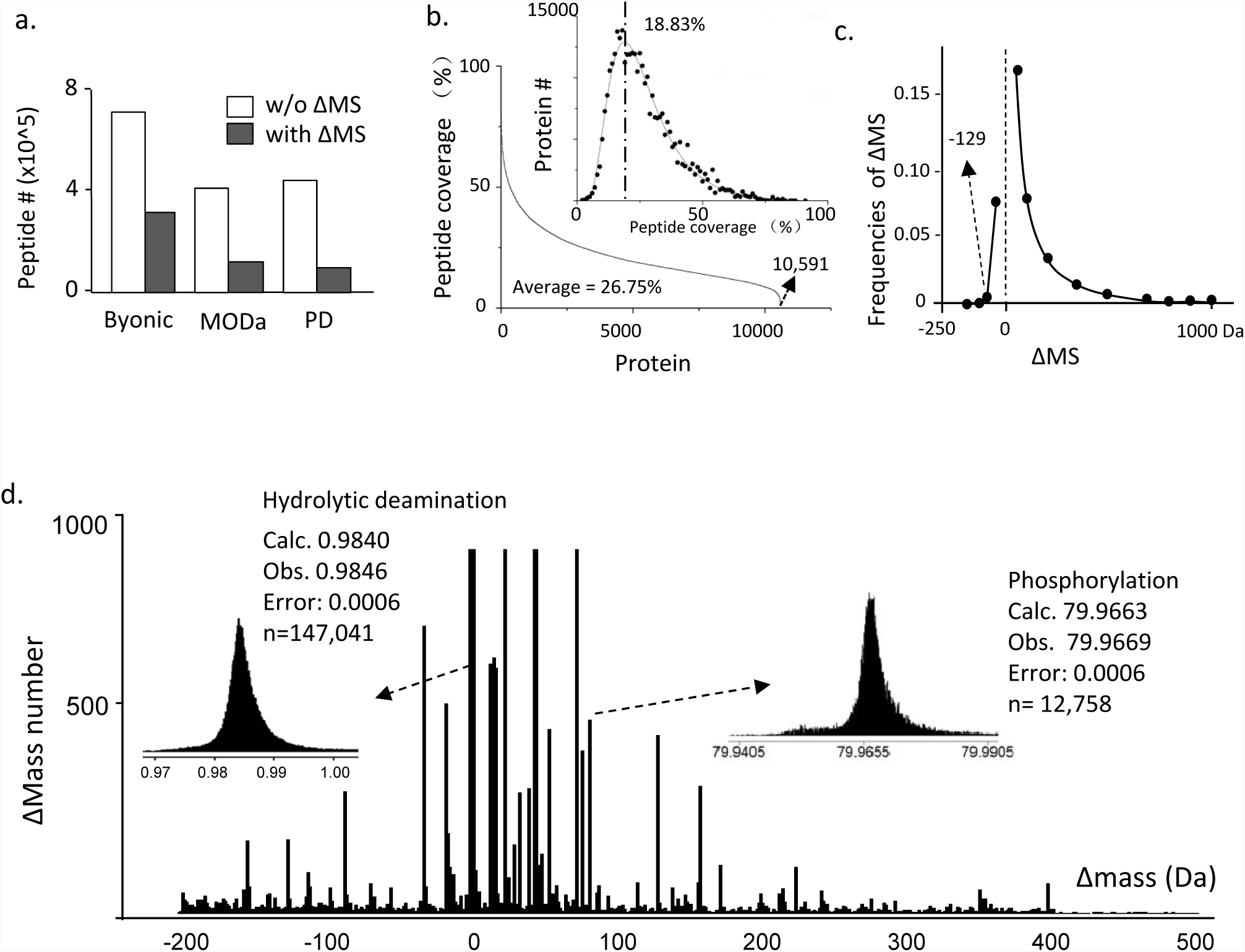
Widespread nonzero delta masses in human spermatozoa. **a**, The number of peptides with (solid bar) and without (open bar) nonzero delta masses (ΔMS) obtained by searching MS/MS datasets with Byonic, MODa, and Protein Discover (PD). **b**, Peptide coverage of the 10,591 non-redundant proteins identified in human spermatozoa. The average peptide coverage was 26.75%. The upper-right panel shows the protein number against peptide coverage (most proteins had a coverage of 18.83%). **c**, Occurrence frequency of Δ masses at low resolution. The smallest mass loss was129 Da, and most of the mass gains were smaller than 500 Da. **d**, Occurrence frequency of Δ masses at high resolution. Nonzero Δ masses from −200 to +500 Da were rounded to four decimal places, and frequencies plotted against Δ masses. The upper-left panel is an enlargement of the Δ mass peak at 0.9838 Da, and the upper-right panel is the Δ mass peak at 79.9669 Da. Calc., calculated; Obs., observed.

Most nonzero delta masses were between −129 to 500 Da, and the occurrence of delta masses with smaller gains and losses increased exponentially (**Fig. 1c**). It should be noted that a delta mass of −129 Da is the theoretical maximum mass loss due to amino acid substitution, equivalent to the replacement of Tryptophan with Glycine. Thus, we reasoned that the mechanism underlying the mass loss of the ncAAs was largely due to amino acid substitutions, assumedly by single nucleotide polymorphisms (SNPs) at the DNA level or other not yet defined mechanisms. On the contrary, most of mass gains were undefined delta masses smaller than 500 Da that did not match amino acid substitutions or previously reported PTMs.

To characterise these delta masses, their frequencies were plotted against delta masses rounded to four decimal places. Indeed, delta masses were highly clustered (**Fig. 1d**), and some accurately matched previously reported protein modifications or amino acid substitutions. Besides that matching oxidation, the most abundant peak was 0.9846 Da, which matches hydrolytic deamination (0.9840 Da, error = 0.0006, n = 147,041). A well-documented peak was 79.9669 Da, which matches a phosphorylation modification (79.9663 Da, error = 0.0006, n = 12,758). In general, these delta masses with previously reported PTMs and amino acid substitutions, such as phosphorylation, acetylation, methylation, and ubiquitylation, were usually identified with high accuracy and high occurrence frequency. However, most of the clustered delta masses, especially those with sharp peaks and high abundance, were unknown and considered as undefined PTMs, amino acid substitutions, or generally ncAAs.

### Multivariate Gaussian clustering and machine learning determination of high-confidence ncAAs

Based on clustering, these delta masses corresponded to molecular differences of some known and mostly unknown ncAAs. The widths of clustered peaks were typically within the lower ppm range and hence compatible with the accuracy of the Orbitrap mass spectrometer. Given that errors were random, clusters with unknown delta masses were assumed to reflect novel protein modifications, substitutions or generally ncAAs. To quantitatively identify all ncAAs, delta masses were divided into subgroups with 1 Da intervals and analysed by multivariate clustering using Gaussian mixture components (Fraley et al., 2012). Gaussian regression was followed to each cluster to determine the peak value (expected cluster), the standard deviation (SD), and the Goodness-of-Fit (R^2^), which resulted in 617 qualified delta masses (**Supplemental Table 2**). These delta mass clusters were further compared with the delta masses previously reported PTMs and substitutions, including those found in the UniMod (Creasy and Cottrell, 2004), RESID (Garavelli, 2004), ExPASy (Gasteiger et al., 2003) and ABRF (Arnott et al., 2003) databases (**Supplemental Table 3**). As a result, 192 clusters closely matched the previously reported delta masses, and 232 did not. In total, 424 clustered delta masses were confidently assigned (**Supplemental Table S4**). Among the unmatched clusters, 136 were qualified by R^2^ >0.2, and 96 reflected selective occurrence over a given amino acid (frequency >50%). It should be noted that the number of matched clusters did not significantly vary with R^2^ value (**Fig. 2a**); however, the number of unmatched clusters dramatically increased when R^2^ <0.2, representing a turning point for unreliable clusters.

**Fig. 2.**
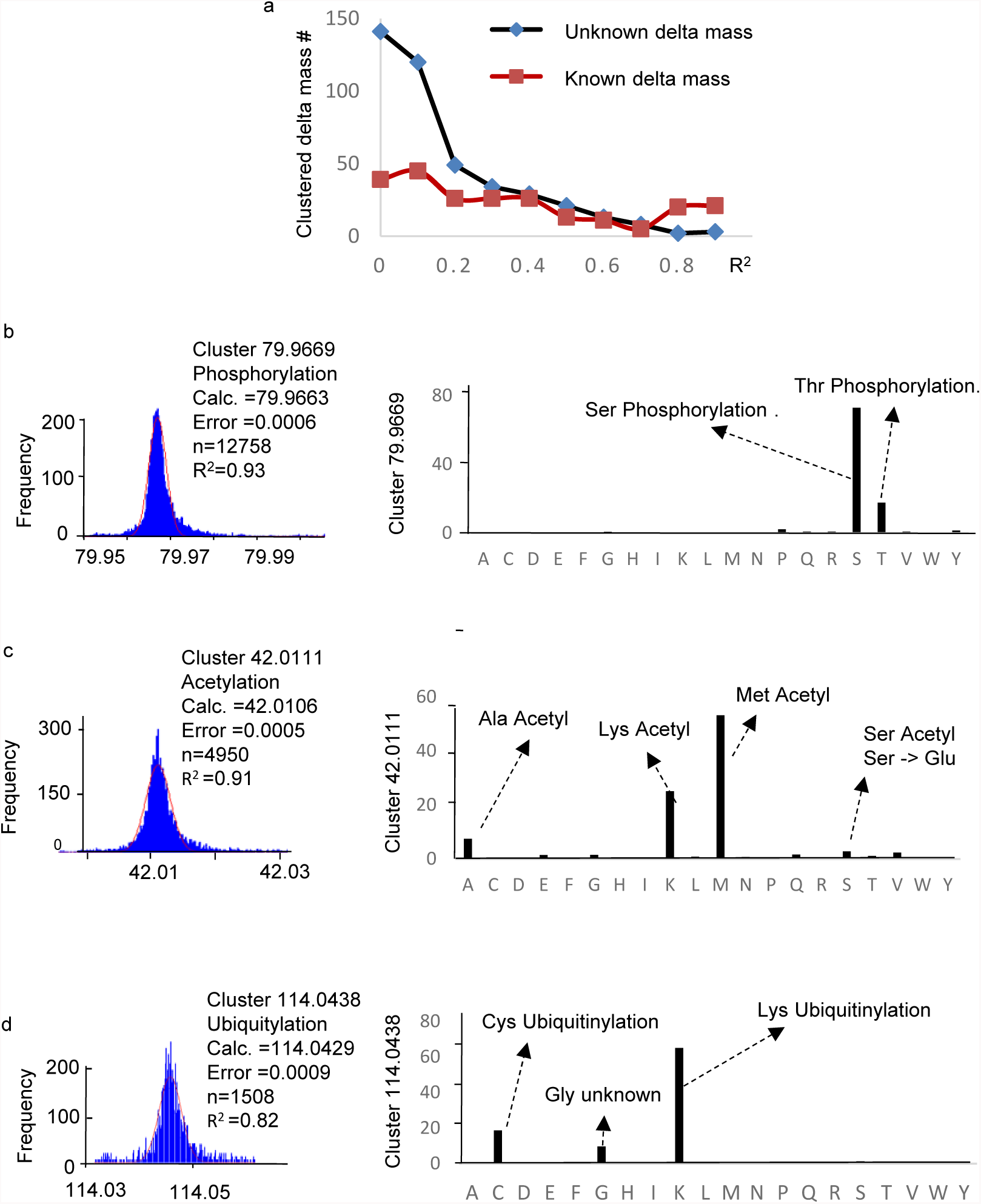
Multivariate clustering and Gaussian regression of nonzero delta masses. **a**, The number of clustered delta masses from matched (square) and unmatched (diamond) groups correlated with the Goodness-of-fit (R^2^) of the clusters. Matched clusters were positively identified even with low R^2^ values, whereas unmatched clusters were considered unreliable when R^2^ <0.2. **b**, Occurrence frequencies of delta masses at each coding amino acid position. Note that oxidation was omitted from the calculation. **c**, Gaussian regression of phosphorylation clustered at delta mass 79.9669 Da (red curve, n = 12758, R^2^ = 0.93), 73% of which were found at Ser and 8% at Thr. However, Tyr phosphorylation only accounted for ~1%. **d**, Gaussian regression of acetylation clustered at delta mass 42.0111 Da (red curve, n = 4950, R^2^ = 0.91), with 53% found at Met, 25% at Lys and 7% at Ala. **e**, Gaussian regression of ubiquitylation clustered at delta mass 114.0438 Da (red curve, n = 1508, R^2^ = 0.82), with 65% found at Lys, 19% at Cys and 7% at Gly.

Notably, these clustered delta masses were detectable from almost all standard amino acids with a certain level of false positive occurrence (**Supplemental Table S4**). However, those amino acids with previously reported delta masses were indeed found with high occurrence frequencies and matched with known PTMs and/or substitutions (**Supplemental Table 3**). They were considered true positive ncAAs. For example, the cluster of 79.9669 ± 0.0019 Da (n = 12,758, R^2^ = 0.93) matched phosphorylation (**Fig. 2b**, left panel), which is reported at Ser, Thr, and Tyr and important for signal transduction (Hunter, 1995). In human spermatozoa, two true positive ncAAs at Ser+79.9669 (73%) and Thr+79.9669 (18%) were identified (**Fig. 2b**, right panel). Similarly, delta masses clustered at 42.0111 ± 0.0019 Da (n= 4950, R^2^ = 0.91) matched acetylation (**Fig. 2c**, left panel), which is a modification of the epsilon-NH3 of Lys and alpha-NH3 of the N-terminal amino acid (Grunstein, 1997; Liu et al., 2008; Yi et al., 2011). In human spermatozoa, at least three true positive ncAAs including Lys+42.0111 (25%), Met+42.0111 (53%), and Ala+42.0111 (7.5%) were identified (**Fig. 2c**, right panel). Another cluster, 114.0438 Da (n = 1508, R^2^ = 0.82), matched ubiquitylation (**Fig. 2d**, left panel), which was detected with high frequency for all three true ncAAs including Lys+114.0438 (65%), Cys+114.0438 (19%) and Gly+114.0438 (10%; **Fig. 2d**, right panel). Ubiquitylation at Lys and Cys has been previously reported (Bhoj and Chen, 2009; Hirsch et al., 2009; Sakamoto et al., 1999), whereas Gly+114.0429 Da has not been reported previously. Finally, the cluster at −45.9872 (n = 736, R^2^ = 0.74) matched Cys-Gly substitution, and was detected from one dominant true positive ncAA C-45.9872 (91%). In summary, from the 192 matched delta mass clusters, we identified 277 ncAAs, among which 137 were true possives that successfully matched with previously reported PTMs/substitutions (**Supplemental Table 3**) and 140 were unreported but occurred with high frequency and/or corresponded to chemically reactive modifications or substitutions.

To identify ncAAs for the 232 unmatched delta mass clusters, the ncAA results of the matched delta masses were used as a training dataset to build a decision tree model (Smith and Frank, 2016). Additional criteria were as follows: (1) each known cluster was typically reacted with not more than four amino acids; and (2) the total occurrence frequency of high-confidence ncAAs for each cluster was greater than 80%. The sensitivity and specificity of the model were 94.9% and 98.1%, respectively, for the matched delta masses. By applying this decision tree model to the unmatched delta mass clusters, we predicted 294 ncAAs with high sensitivity and specificity. Therefore, a total of 571 ncAAs were identified in human spermatozoa. Among them, 137 were previously kown PTMs and/or substitutions, and 434 were not previously reported (**Supplemental Table 5**).

### Numerous novel noncoded amino acids with undefined side-chain structures and unreported mechanisms

Remarkably, a large number of delta masses and ncAAs did not match previously known protein modifications and substitutions (**Fig. 3a**). These unreported delta masses were reasoned to form novel ncAAs with hitherto undefined side-chain structures and unreported mechanisms. However, many of the delta masses were highly selective and dependent on the chemical properties of the targeted amino acids, indicating putative modification mechanisms. For instance, some unreported delta masses occurred more frequently on basic residues, including delta mass 211.0968 ± 0.0036 Da (n = 1895, R^2^ = 0.62) on Lys (77%) and Arg (22%), and delta mass 353.2445 ± 0.0028 Da (n = 948, R^2^ = 0.58) on Lys (25%) and Arg (72%). Meanwhile, other unreported delta masses were selective for acidic residues, such as delta mass 52.9126 ± 0.0028 Da (n = 3342, R^2^ = 0.89) on Asp (23%) and Glu (22%). These results indicated a preference for reacting with the carboxyl group of acidic amino acids. In addition, the abundant delta mass cluster at 223.0347 ± 0.0028 Da (n = 3569, R^2^ = 0.91) was predominantly associated with the Cys residue (93%), indicating preferential reactivity with a thiol group. Thus, although the chemical structures were unknown, the selectivity of these ncAAs is indicative of their unique reactivity with the side-chains of the modified amino acids.

**Fig. 3.**
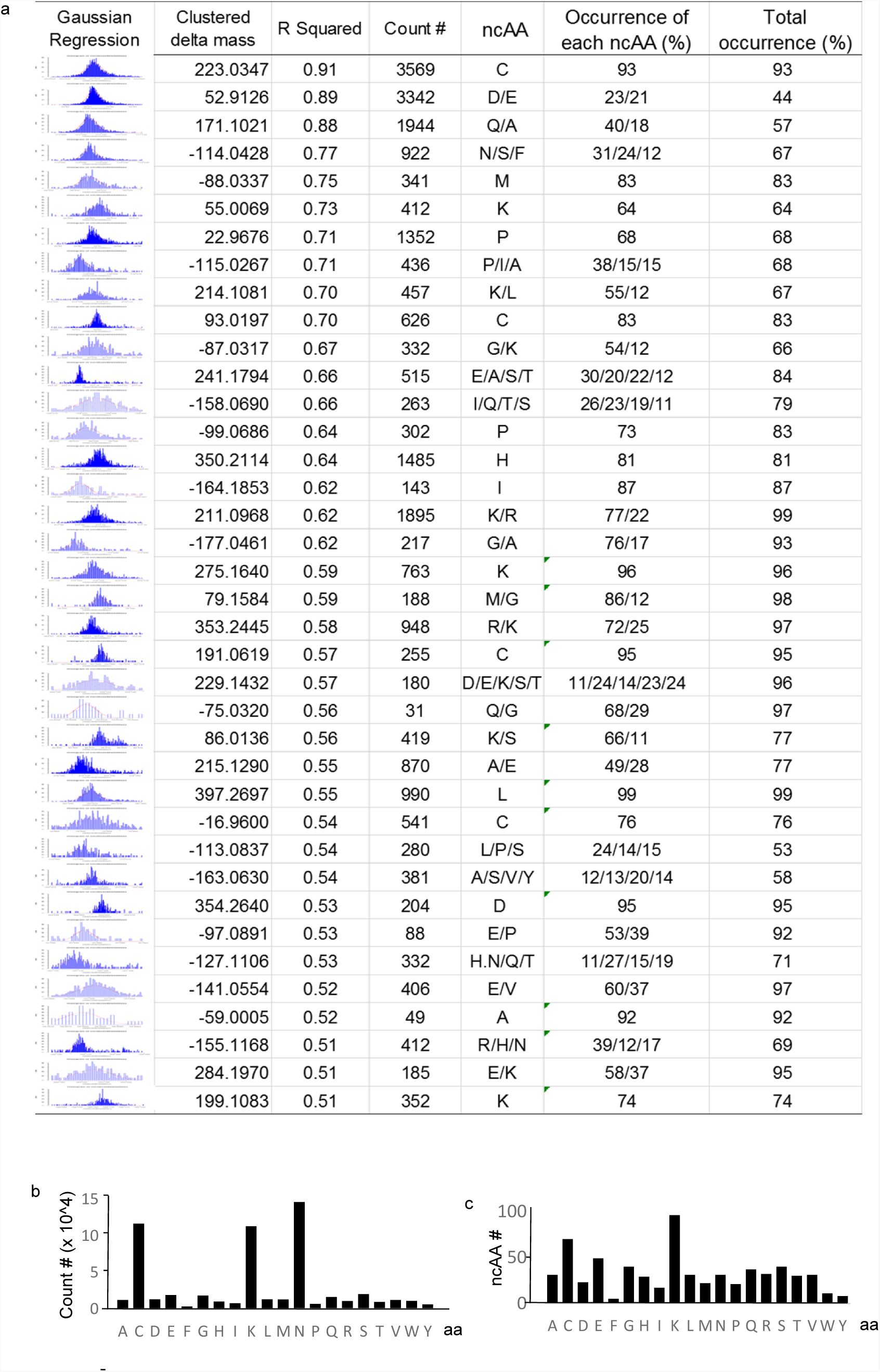
Machine learning prediction of novel ncAAs with undefined side-chain structures. **a,** The most abundant delta masses and ncAAs not previously reported. Blue bars, detection of delta masses. Red curve, Gaussian regression (R^2^ >0.5, count # >50, occurrence >10%). **b,** Total occurrence number of delta masses on standard amino acids. Gln (n = 14,600), Cys (n = 11,700) and Lys (n = 11,300). **C,** Lys was the most reactive residue, targeted by 96 different ncAAs, compared with 70 Cys, 40 for Ser and 40 for Gly.

In summary, the most vulnerable amino acids for ncAAs in sperm proteins were Asn (n = 146,035, 25%), Cys (n = 116,520, 20%) and Lys (n = 113,032, 19%), which accounted for 65% of all delta masses (**Fig. 3b**). However, the most reactive amino acids for ncAA were Lys, Cys and Glu, which were targeted by 96, 70 and 49 different mutations/substitutions, respectively (**Fig. 3c**).

### Widespread and polymorphous ncAA-containing protein sites

For 571 ncAAs, a total of 29,053 non-redundant ncAA-containing protein sites were identified in human spermatozoa (**Supplemental Table 5)**. With a 26.75% protein coverage, the actual number of ncAA-containing protein sites was estimated to be up to 108,609, indicating the widespread existence of ncAA protein sites that might have a potentially crucial impact on protein structure and function. Interestingly, the proteins most often targeted by ncAAs were A-kinase anchor protein 3/4 (AKAP3/4, n = 33,705), lactotransferrin (TRFL, n = 23,004), semenogelin-I/II (SEMG-I/II, n = 39,798), keratin type II cytoskeletal 1 (K2C1, n = 8378; **Fig. 4a**). Some proteins were highly reactive to various different ncAAs, including SEMG-II, which was targeted by 646 ncAAs, AKAP4, which was targeted by 610 ncAAs, and TRFL, which was targeted by 538 ncAAs (**Fig. 4b**). Taking SEMG-II as an example, 646 ncAAs spanned 267 amino acid positions, suggesting multiple ncAAs had to compete for limited amino acid positions. Among them, eight different ncAAs were found at Gln134 of SEMG-II (**Fig. 4c**), including Q+12.00045 (unknown, n = 32), Q-57.02116 (Gln>Ala substitution, n = 15), and Q+128.09574 (unknown, n = 12). Meanwhile, 47 different ncAAs were identified at SEMG-II Cys159, including C-33.9871 (dehydroalanine, n = 1091), C+72.02261 (unknown, n = 421), C+47.98546 (oxidised to cysteic acid, n = 370), and C+75.99925 (mercaptoethanol, n = 291).

**Fig. 4.**
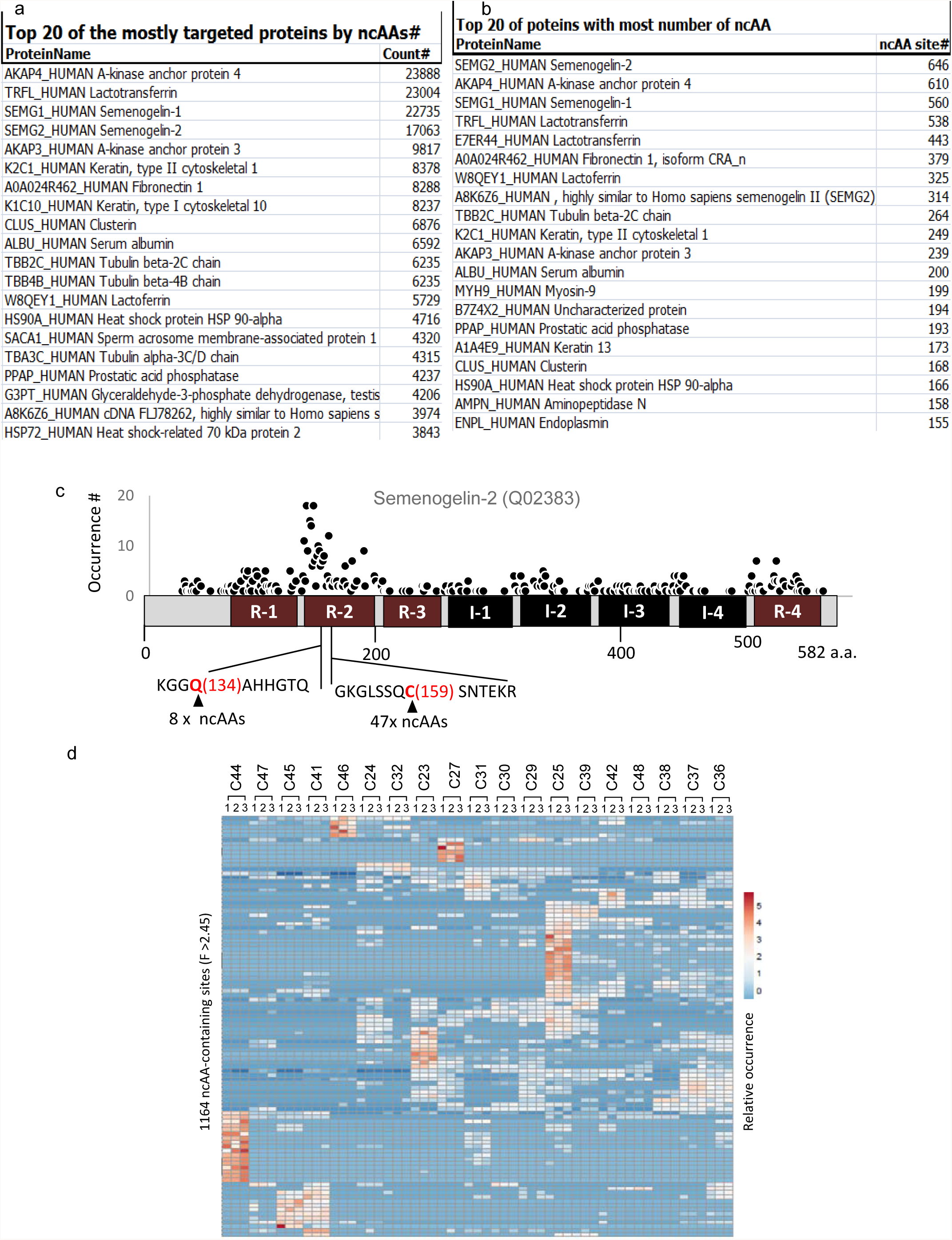
Polymorphous ncAA proteins and ncAA protein sites. **a,** The 20 proteins with the most ncAA occurrence. **b,** The 20 proteins with the highest number of different ncAAs. **c**, Occurrence and protein sites for semenogelin-2 (SEMG-II, Q02383). R-1 to R-4, the repeat region with unidentified features. I-1 to I-4, type I tandem repeats. KGG**Q**(134)AHHGTQ, Q134 was targeted by 18 different ncAAs. GKGLSSQ**C**(159) SNTEKR, C159 was targeted by 35 different ncAAs. **d,** Clustering analysis of ncAA-containing protein sites. Spermatozoa samples from 19 healthy people (C21-C47) was performed in triplicate (lane 1-3) and ncAA protein sites (F >2.45, p <0.01) were clustered with Principal Component Analysis (Metsalu and Vilo, 2015).

To analyse the polymorphous nature of ncAAs, variations between healthy human subjects were analysed using ANOVA, and 351 ncAA-containing protein sites varied significantly between the healthy individuals (F >F_0.01_ (18,38) = 2.45, p <0.01, n>20; **Supplemental Table 6**). When clustered using Principal Component Analysis (Shen et al., 2015), variations were largely the result of a significant increase in the occurrence of ncAAs in one or a few individuals (**Fig. 4d)**, implying a polymorphous nature for ncAAs in healthy populations. The most abundant polymorphism was C-33.9871 (dehydroalanine, F = 3004), which was only detected at Cys12 of the tubulin beta-2C chain (TBB2C) in individual C27. Another example was E+17.04763 (unknown, F = 241), which was only detected at E322 of meiosis-specific nuclear structural protein 1 (MNS1) in individuals C44 and C31. In addition, D-15.9733 (unknown, F = 477) was only detected at D353 of annexin A11 (ANX11) in individual C44. These results suggest that polymorphous ncAA-containing protein sites contribute to population diversity and even human diseases.

### Discrimination of patients with oligoasthenospermia by ncAAs

The clustered delta masses and ncAAs were compared between healthy humans and patients with oligo-asthenospermia, a male reproductive disease causing low sperm count and poor sperm motility. At the delta mass level, 62 of the identified clusters were up- or down-regulated > 2-fold in oligoasthenospermia patients (**Fig. 5a; Supplemental Table 7**). The most abundant down-regulated delta masses were 79.9668 (phosphorylation, −2.3-fold), 17.9567 (Leu/Ile>Met substitution, −3.67) and 211.0968 (unknown, −3.2-fold), whereas the mostly abundant up-regulated was 344.1659 (unknown, 18.0-fold). At the ncAA level, 58 ncAAs (10%) were up- or down-regulated >2-fold (**Fig. 5b**; **Supplemental Table 8**). Consistently, these included S+79.9668 (Ser phosphorylation, −2.3- fold), T+79.9668 (Thr-phosphorylation, −2.5-fold), K+211.09682 (unknown, - 3.2-fold), K+42.0111 (acetylation, −6.4-fold) and E-14.0132 (E>D, +2.9-fold). These results demonstrated that many delta masses and their ncAAs were dysregulated in oligoasthenospermia, most of which were not previously reported. These oligoasthenospermia-associated ncAAs presumably alter the side-chain structures and, ultimately, the overall protein function, thus contributing to pathogenesis.

**Fig. 5.**
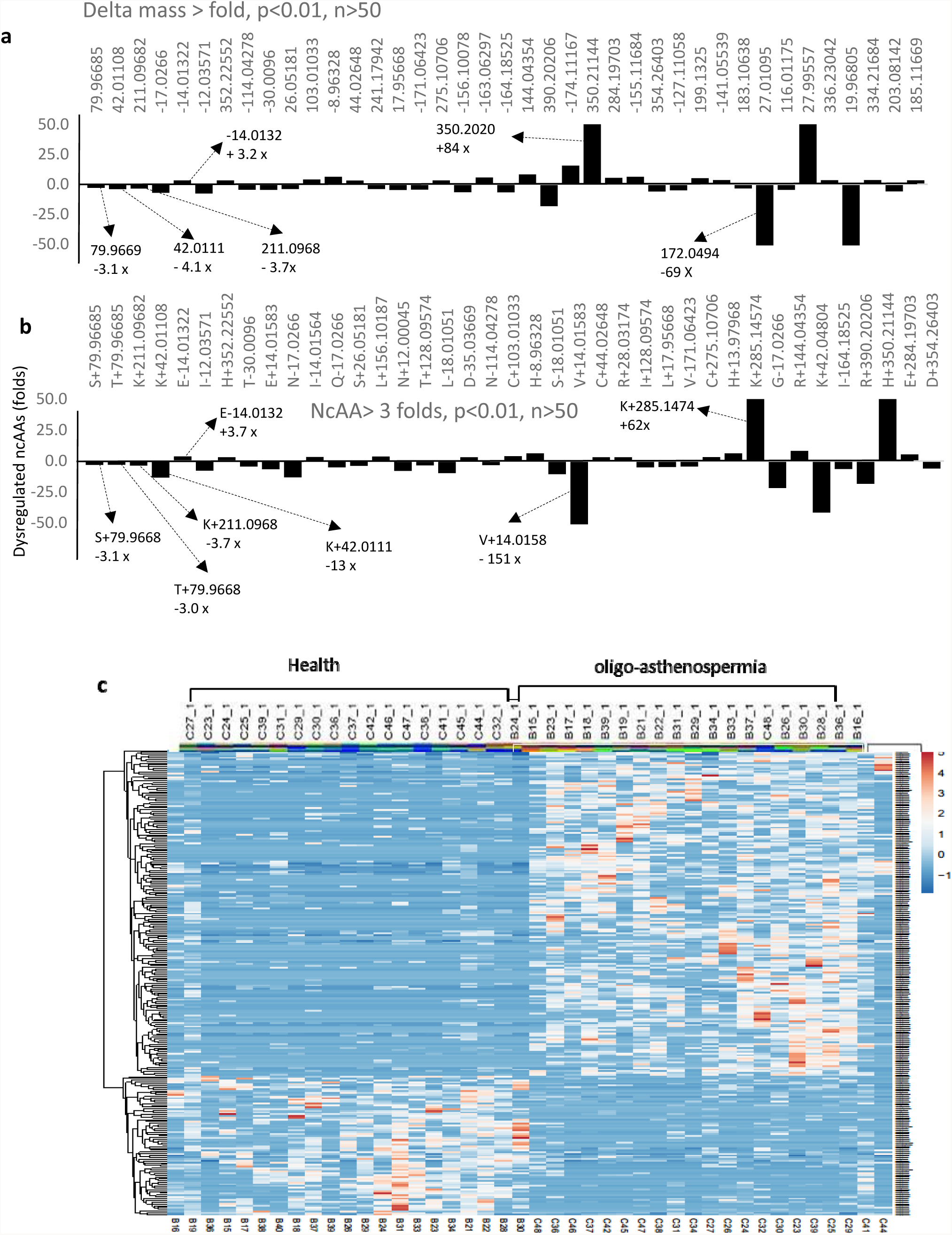
Discrimination of patients with oligoasthenospermia by delta masses and ncAAs. **a,** Up/down-regulated delta masses in oligoasthenospermia. Of 424 clustered delta masses, 62 were up- or down-regulated >3-fold in patients (R^2^ >0.2, p <0.01, n >20). **b,** Up/down-regulated ncAAs in patients. Of 571 ncAAs, 58 were up- or down-regulated >3-fold (R^2^ >0.2, p <0.01, n >20). **c,** Clustering analysis of up/down-regulated ncAA-containing protein sites. Of 29,053 ncAA-containing protein sites, 332 were discriminated into healthy and diseased groups (ratio >2, R^2^ >0.2, p <0.01, n >20). Upper-right panel, up-regulated ncAA protein sites associated with oligoasthenospermia. Lower-left panel, ncAA protein sites down-regulated in patients but presumably required by healthy people.

To provide further support, ncAA-containing protein sites were clustered by Principal Component Analysis (Shen et al., 2015), which clearly discriminated results into two distinct panels (**Fig. 5c**). The lower-left panel includes the up-regulated ncAA protein sites associated with oligoasthenospermia, whereas the upper-right panel includes ncAA protein sites that are characteristic of healthy individuals. In total, 332 out of 29,053 ncAA-containing protein sites were up- or down-regulated >2-fold (**Supplemental Table 9**). The most abundant and highly regulated protein sites are shown in **Fig. 6a**. Interestingly, SEMG and AKAP families were reciprocally regulated: all ncAA-containing sites of SEMGs were up-regulated in oligoasthenospermia (**Fig. 6b**), whereas most ncAA sites of AKAPs were down-regulated (**Fig. 6c**). A typical example was Ser- and Thr-phosphorylation, which were exclusively down-regulated in AKAP4. Notably, most of the oligoasthenospermia-associated ncAA sites have not been previously reported. For example, the C-2.0147 modification at Cys908 of the coiled-coil domain-containing protein (Q96JN2) occurred 108 times in healthy people, but only twice in oligoasthenospermia patients (**Supplemental Table 9**), suggesting a significant loss of disulfide bonds in the acrosome during spermatogenesis and fertilisation (Geng et al., 2016). In conclusion, our results demonstrated that many ncAA protein sites were associated with oligoasthenospermia, implicating them in pathogenesis and suggesting they might be potential biomarkers for diagnosis and targeting.

**Fig. 6.**
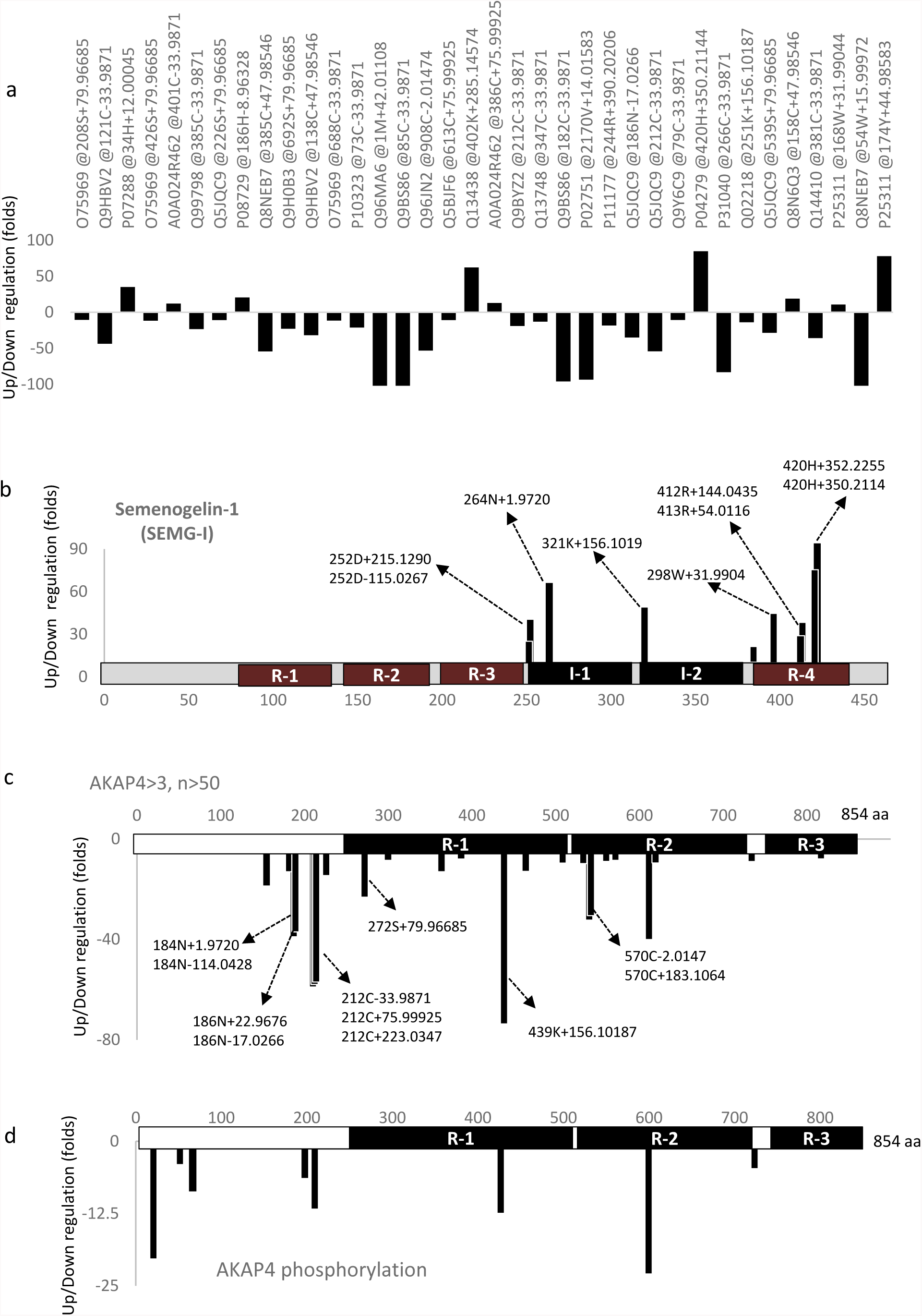
Oligoasthenospermia-associated ncAA proteins and ncAA protein sites. **a,** The most abundant and regulated ncAA-containing protein sites. The ncAA protein sites were assigned by protein ID followed by amino acid position and ncAA. Positive/negative, up-/down-regulated ncAA protein sites (ncAA protein sites with p <0.01, n >50, and up/down-regulation >10-folds are shown). **b**, The ncAA-containing sites in semenogelin-1 (SEMG-I) up-regulated in oligoasthenospermia patients. The ncAA-containing sites were assigned by amino acid position followed by ncAA. R-1 to R-4, the repeat region with unidentified features. I-1 to I-4, type I tandem repeats. **c,** The ncAA sites of A-kinase anchor protein 4 (AKAP4) down-regulated in oligoasthenospermia patients. R-1, R-2 and R-3: the repeat region with unidentified features. **d.** AKAP4 sites at which phosphorylation was decreased in oligoasthenospermia patients

### Large numbers of amino acid substitutions irrelevant to SNPs

Amino acid substitutions among ncAAs were identified by matching the delta masses between the 20 standard amino acids (**Supplemental Table 10**). Among 571 ncAAs, 98 matched with amino acid substitutions spanned 4882 different protein sites (**Fig. 7a**; **Supplemental Table 11**). Among these substitutions, N>D (2932 sites, n = 103,327) and Q>E (330 sites, n = 1804) were the most abundant substitution that share the same delta mass with hydrolytic deamination. The second most abundant class of substitutions occurred between side-chains with similar biochemical properties and reactivity. These included substitutions between Val and Leu/IIe hydrophobic side-chains (48 sites, n = 3647), between Glu and Asp negatively-charged carboxyl-containing side-chains (16 sites, n = 1326), and between Ser and Thr hydroxyl-containing side-chains (39 sites, n = 134). It should be emphasised that many high-frequency ncAA substitutions would require two spontaneous point mutations if they were generated at the genomic level. Examples include G>Q (82 sites, n = 363), A>V (24 sites, n = 117), V>T (6 sites, n = 120) and Q>A (10 sites, n = 157). As the frequency for spontaneous double point mutations was very low, this hints at the existence of as yet undefined mechanisms in addition to SNPs for these ncAA substitutions in humans.

**Fig. 7.**
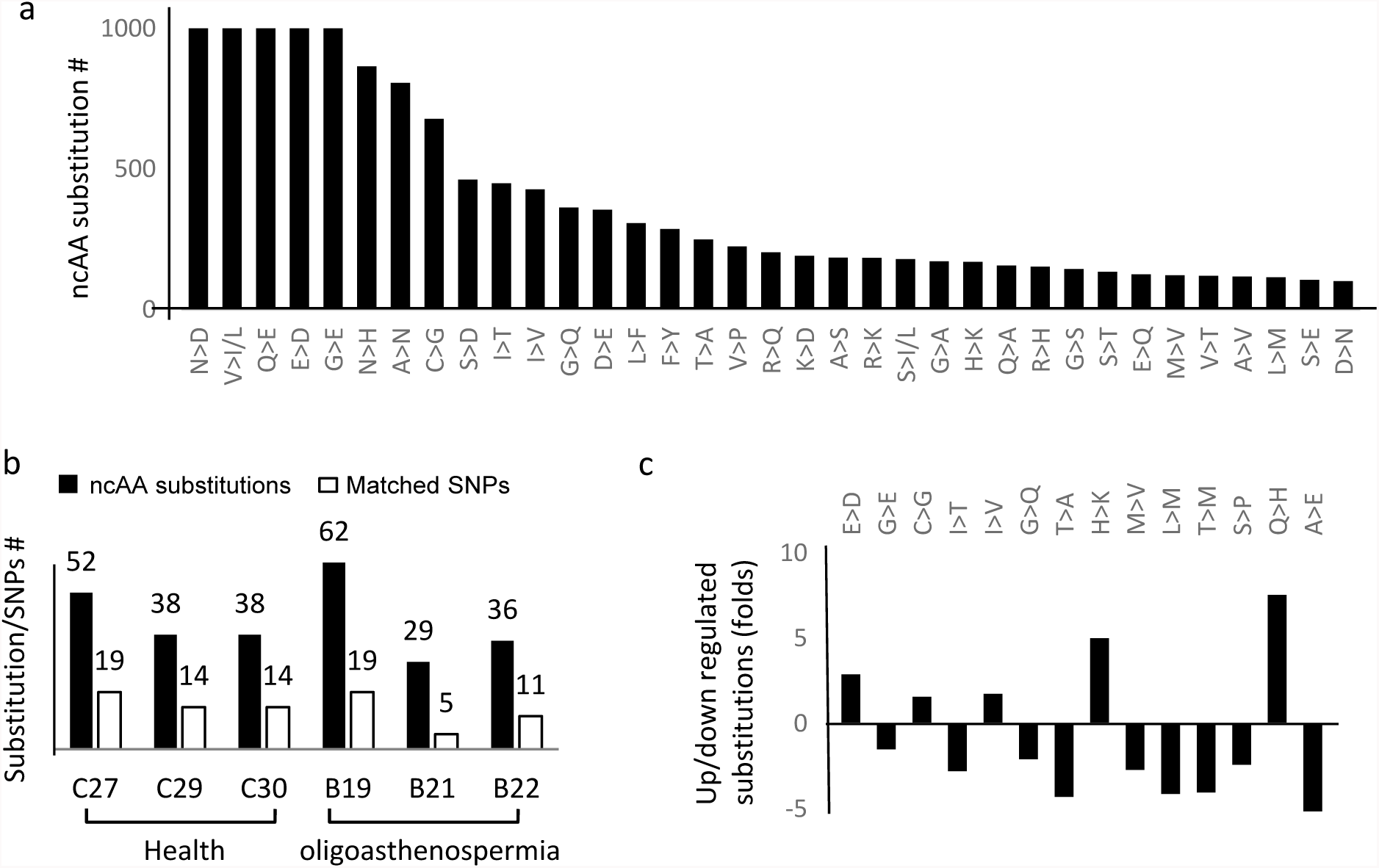
ncAAs matching amino acid substitutions in sperm. **a,** ncAAs were matched with all possible substitutions of the 20 standard amino acids (**Supplemental Table S11a**). The top 35 most abundant ncAA substitutions (n > 100) are shown. **b**, Validation of ncAA substitutions by exon sequencing (**Supplemental Table S13**). Missense SNPs in three healthy individuals and three oligoasthenospermia patients were determined by exon sequencing and compared with ncAA substitutions (solid bars). The ncAA substitutions that matched with missense SNPs are shown in open bars. **c,** oligoasthenospermia-associated substitutions (**Supplemental Table S11b**). The most up-/down-regulated ncAA substitutions (p<0.05, t-test) are shown.

To validate the identified ncAAs in human sperm, ncAA substitutions were determined by exon sequencing. Genomic DNA from three patients and three healthy people was prepared and exon sequences were determined. Missense SNPs were identified (**Supplemental Table 12**) to compare with amino acid substitutions in human sperm ncAAs (**Fig. 7b**). The results demonstrated that 40 out of 98 ncAA substitutions and 103 out of 4882 ncAA substitution protein sites were indeed matched with the missense SNPs (**Supplemental Table 13**), and thus genomic mutations were confirmed as the cause in these cases. However, most ncAA substitutions did not find corresponding SNPs, suggesting additional mechanisms operating at the level of post-transcription, RNA editing, and PTM were responsible for these ncAA substitutions.

In addition, the ncAA-substitution sites also differed between healthy individuals and oligoasthenospermia patients (**Fig. 7c**). The most highly regulated substitutions included G>Q (−23-fold, n = 75), H>K (+20-fold, n = - 136), and I>T (−13-fold, n = 63). The most highly up-regulated site occurred at G239V of carcinoembryonic antigen-related cell adhesion molecule 6 (CEACAM6, +47-fold, n = 46), and the most highly down-regulated sites occurred at A1059V of EF-hand calcium-binding domain-containing protein 6 (EFCAB6, −70-fold, n = 38) and G98Q of sperm mitochondrial-associated cysteine-rich protein (SMCP, −23-fold, n = 75; **Supplemental Table 13**).

## DISCUSSION

Herein, we developed a systematic workflow for identifying all possible protein residues not directly encoded by their genomic sequences, namely noncoding amino acids (ncAAs), in proteome. By measuring the delta masses between actual protein residues and coding amino acids, over a million nonzero delta masses were detected in human spermatozoa. These delta masses were grouped into 424 high-quality Gaussian clusters, and 571 mostly novel ncAAs spanning 29,053 protein sites were identified with confidence, suggesting a total of over 108,000 ncAA protein sites in humans. According to the definition, these ncAAs would cover the non-standard protein residues generated by point mutations and genomic editing (Drezen et al., 2016; Tak and Farnham, 2015), RNA editing and transcriptional mutagenesis (Bregeon and Doetsch, 2011; Yang et al., 1995), PTMs, and erroneous protein synthesis (Beltrao et al., 2012; Drummond and Wilke, 2009; Mathias et al., 2015). Therefore, this proteomic approach identifies known and unknown PTMs, amino acid substitutions, missense SNPs, and all possible ncAAs with yet undefined mechanisms.

Strikingly, most of the delta masses and ncAAs corresponded to unresolved side-chain structures and were not previously reported. These novel ncAAs were chemically different from the standard amino acids and have different primary and presumably tertiary structures, ultimately affecting protein function. By searching known delta mass databases, common PTMs were confidently identified and well covered, indicating the reliability of the approach. Well-known PTMs such as phosphorylation, acetylation, methylation and ubiquitination were the most abundant of the identified ncAAs. The reliability of the unmatched delta mass clusters and novel ncAAs were strongly supported not only by their high frequencies and sharp peaks assigned by Goodness-of-Fitness of Gaussian regression with confidence, but also due to their high selectivity of occurrence over unique coding amino acids. Some of the unknown ncAAs occurred as frequently as acetylation and methylation, such as E/D+52.9126 (n = 3342, R^2^ = 0.89), K+55.0069 (n = 412, R^2^ = 0.73), and P+22.9676 (n = 1352, R^2^ = 0.71), suggesting that their importance for protein structure and function. Interestingly, even for known delta masses, many of their ncAA-containing proteins and/or ncAA sites were not previously reported. The peptide coverage of the 10,591 non-redundant proteins was only 26.75%, implying the presence of over 108.609 known and unknown ncAA protein sites in human spermatozoa. Therefore, our approach revealed the widespread existence of unknown ncAAs with yet undefined side-chain structures and widespread ncAA-containing protein sites, representing a significant advance in our understanding of protein structure and function.

The polymorphous nature of ncAAs was supported by significant variations between different individuals, suggesting that ncAAs contribute to amino acid polymorphisms at the protein level and important for the diversity of human populations. Notably, ncAA-caused amino acid polymorphisms at the protein level have 571 dimensions of freedom, whereas nucleotide polymorphisms at the DNA level vary through four nucleotides. Thus, ncAAs provide far more complex of regulations on protein structure and function. Particularly, protein amino acid polymorphisms by novel ncAAs usually indicate undefined enzymatic reactions on at the side-chains of the identified ncAA protein sites in a subpopulation.

Importantly, ncAAs were demonstrated to discriminate human diseases. Significant variations were observed between healthy individuals and patients with oligoasthenospermia in terms of delta mass clusters, ncAAs and ncAA-containing protein sites. Unlike SNPs, disease-associated ncAAs were considered relevant to specific biochemical reactions, and likely cause protein malfunctions, abnormal signal transduction, and pathogenic alteration in human diseases. Thus, the disease-associated ncAAs and ncAA-containing protein sites might be developed as diagnostic biomarkers and therapeutic targets.

By comparing with the standard amino acids, ncAAs in human spermatozoa were classified into amino acid substitutions and chemical modifications. Approximately 13% of ncAAs were matched with amino acid substitutions, among which 40% were due to genomic mutations (SNPs), as confirmed by exon sequencing. It validated the efficiency of the ncAA approach that the ncAA substitutions corresponded with missense SNPs with consistence between the DNA and protein levels. Notably, 60% of ncAA substitutions did not match missense SNPs, leading to the hypothesis that most ncAA substitutions were generated by mechanisms additional to genomic point mutations, such as post-transcriptional RNA editing (Yang et al., 1995), erroneous protein synthesis (Ribas de Pouplana, 2014) (Beltrao et al., 2012; Drummond and Wilke, 2009; Mathias et al., 2015), and/or protein editing (Shen et al., 2015).

In summary, widespread ncAAs were identified in human spermatozoa using a systematic approach, most of which were unreported PTMs and amino acid substitutions. These ncAAs were polymorphous and discriminative in healthy and diseased populations. This approach could be applied to different human tissues and organs to systematically identify disease-associated ncAAs that are potential biomarkers for diagnosis and drug targeting. The method opens up a new dimension for understanding protein structure, function, and regulation relevant to physiological and pathological mechanisms.

## Author Contributions

J.-H.Y. designed the project and wrote the manuscript, X.C., J.G., X.K., M.S. Y.W. and X.W. conducted data mining, X.L., F.W., F.S., and S.S. acquired the data, C.L., B.B., W.H., T.H., H.Z. and H.Z. provided sample collection and preparation, J.-H.Y., Y.E.C., K.M.A., Z.C., and Z.-J.C. provided interpretation and discussion.

## Acknowledgements

This work was supported by grants from the National Natural Science Foundation of China (31471322, 30928031, 81430029, 81490743, 81622021, 81601256, 31371453, and 31571548), the National Research and Development Plan (2016YFC1000604), and Shandong Science & Technology Bureau. The work was partially supported with resources and the use of facilities at VA Boston Healthcare System, USA, Shandong University Cancer Research Center and Reproductive Medical Center, Shandong Computer Science Institute, Xian Fourth Military Medical University, and Shandong Food and Drug Research Institute. We declare that we have no financial and personal conflict of interest.

## EXPERIMENTAL PROCEDURES

### Sample preparation

Human sperm samples were collected from 24 healthy people and 24 patients with asthenospermia under the protocol approved by the Institutional Review Board of Reproductive Medicine, Shandong University. Written consent was obtained from all participants. Sperm samples were washed separately three times with ice-cold PBS (0.1 M Na2HPO4, 0.15 M NaCl, pH 7.5), and re-suspended in 1 ml of chilled 1× RIPA lysis buffer (Millipore 20-188, USA) supplemented with 10 μl of 500 mM DTT and 10 μl of protease inhibitor cocktail (Roche, Switzerland) to protect against oxidation and degradation. After sonication three times for 5 s on ice, lysates were centrifuged at 13,000g for 20 min at 4°C. Supernatants were collected and protein concentrations determined using the Bicinchoninic Acid (BCA) assay (Beyond, China).

### Pre-separation and trypsin digestion

One hundred micrograms of lysate proteins were resolved by 15% SDS-PAGE, stained with Coomassie Brilliant Blue, and divided into 5-10 fractions based on molecular weight. Gel slices were washed with 50 mM ammonium bicarbonate, dehydrated with acetonitrile, and lyophilised in a SpeedVac (Thermo Scientific). Proteins were digested with in-gel sequencing-grade modified trypsin (Promega) at a protein:trypsin ratio of 25:1 in 50 mM ammonium bicarbonate (pH 8) overnight at 37°C. Peptides in the supernatant were collected on ZipTip C18 columns (Sigma-Aldrich), washed with 0.1% trifluoroacetic acid, and eluted with 50% methanol. Desalted peptides were dried in a SpeedVac for LC-MS/MS analysis.

### LC-MS/MS analysis

Desalted peptides were subjected to peptide fractionation by liquid chromatography on an EASY-nLC 1000 system (Thermo Scientific, San Jose, CA). Samples were trapped on a C18 pre-column and fractionated with a long C18 column (300 × Ø 0.075 mm, 3 μm; Reprosil, Germany) with a 180 min linear gradient of 5-35% acetonitrile/0.1% formic acid at a flow rate of 250 nl/min (solvent A, 0.1% formic acid in water; solvent B, 0.1% formic acid in acetonitrile). MS and MS/MS spectra were acquired by a LTQ-Orbitrap Elite mass spectrometer (Thermo Scientific, San Jose, CA) in data-dependent mode. The spray voltage was 2.1 kV and the capillary temperature 275°C. MS spectra were acquired in profile mode in the m/z range of 350-1800 at a resolution of 60,000 at 400 m/z. The automatic gain control was 1×10^6^. MS/MS fragmentation of the 30 most intense peaks was performed for every full MS scan in the collision-induced dissociation mode. Normalised collision energies were set to 35% with an activation time of 10 ms for the MS/MS method. The max precursor ion injection time for MS/MS was 100 ms. The repeat count was set to 2, and the dynamic exclusion duration was 90 s. The minimal signal required for MS/MS was set at 1000. Eight technical replicates were analysed for each sample. LC-MS/MS data for all 48 sperm samples have been submitted to PRIDE Archive (http://www.ebi.ac.uk/pride/archive/).

### Wide-tolerance searching

MS/MS spectra were searched against the human protein database with wide MS tolerance using MASCOT (Perkins et al., 1999), SEQUEST (Yates, 2015), MODa (Na and Paek, 2015) and Byonic (Bern et al., 2012). For wide-tolerance searches with SEQUEST HT in Proteome Discover 2.0 (ThermoFisher), the precursor ion tolerance (peptide) was set to 500 Da, the fragment ion tolerance (fragment) was 0.6 Da, two missed cleavages by trypsin were allowed, the false discovery rate (FDR) for peptides was 1%, and the threshold for accepted MS/MS spectra was set at high confidence. For wide-tolerance searching with MODa v1.21, peptide tolerance was set to 1 Da, the fragment tolerance was 0.6 Da, BlindMode was 1, the modification was set from −200 to 1000 Da, two missed cleavages by trypsin were allowed, the FDR for peptides was 1%, the threshold score and the probability of the peptides and spectrum match were 30 and 0.95, respectively. For wildcard searching with Byonic v2.6, MS/MS spectra were first searched normally using Byonic v2.6 to create a focused human protein dataset. Wildcard searching was then performed against the focused protein dataset. The precursor ion tolerance was set to 10 ppm, and the fragment ion tolerance was 0.6 Da. Trypsin (full specific cleavage) was specified as the cleavage enzyme and up to two missed cleavages were allowed. The modification masses of wildcard searches were set from −200 to 1000 Da.

### Nonzero delta mass clustering

Typically, MS/MS spectra were searched against the human protein database using the wildcard search in Byonic software (Bern et al., 2012) and filtered with Score ≥300, DeltaModsScore ≥10, FDR2D ≤0.01 and FDR_uniq.2D ≤0.01. All amino acid residues that were different from coding genomic sequences were identified to calculate nonzero delta masses (delta mass >0.01), the mass differences between the actual and theoretical (coding) amino acids. These nonzero delta masses were assigned noncoded amino acids (ncAAs). For occurrence analysis, delta masses were rounded to four decimal places and the number of redundancies was considered as the frequency of the delta mass. For clustering, delta masses were divided into subgroups with 1 Da intervals bounded by n-0.5 and n+0.5 Da (n = −200 to 1000). Delta masses in each mass window were analysed by multivariate clustering using Gaussian mixture components (Fraley et al., 2012) with the following constraints: (1) peak half-width >1 ppm; (2) peak distance >2 peak widths; (3) cluster size >20. Clusters within each window were determined by the Bayesian Information Criterion (BIC) (Fraley and Raftery, 1998), where larger BIC values indicate a stronger model and confidence in the number of clusters. Next, clusters in each window were fitted individually with Gaussian regression to calculate the peak value (expected delta mass), the SD, and the Goodness-of-Fit (R^2^). Delta mass clusters were categorised by matching with previously known PTMs, and amino acid substitutions in UniMod, RESID, ExPASy, and ABRF databases. Matched clusters were considered as the true delta masses in the examined protein samples. Unmatched clusters were assigned with confidence only when the Goodness-of-Fit R^2^ >0.2 and/or the cluster was predominantly associated at a single amino acid (>50%). An open online server for delta mass clustering is available at http://www.crc.sdu.edu.cn/ncAAsis.

### ncAA identification by the machine learning algorithm

The ncAAs for matched delta masses were manually classified into high confidence (H) for previously reported modifications, moderate confidence (M) for unreported but chemically possible modifications, and low confidence (L) for unreported and chemically unlike modifications (**Supplemental Table S14**). Based on this classification, a score *s* was assigned for every amino acid *i (I = 1…20)* in each delta mass cluster *j (I = 1…192).* Thus, the score *s_i,j_* was computed as follows:

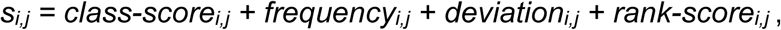

where *class-score_i,j_* is the score transformed from H/M/L classification of amino acid *i* in cluster *j* (*class-score_i,j_ = 2* for class H, *0.5* for M, *0* for L), *frequency_i,j_* is the occurrence frequency of amino acid *i* in cluster *j*,

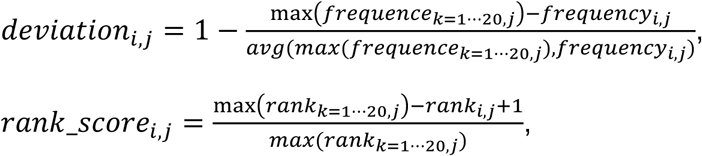

where *rank_i,j_* is the rank of amino acid *i* in cluster *j ordered by* occurrence frequency (parallel ranking available). Using this scoring function, all amino acids were scored and those with s ≥ 2.3 were defined as ncAAs. As the H/M/L classification for amino acids of unmatched delta mass clusters was missing, the scoring method was derived using a decision tree model trained on the dataset of all amino acid occurrences from matched delta mass clusters. In more detail, a five-dimensional vector was derived for every amino acid, with the following elements: (i) *frequency*_*i*,*j*_, (ii) *deviation*_*i*,*j*_, (iii) *rank_score*_*i*,*j*_, and (iv) ncAA or not. The decision tree was trained using J48 in the WEKA software (Smith and Frank, 2016) and evaluated by 8-fold cross-validation.

### Whole exome sequencing

To validate amino acid substitutions, human genomic DNA was extracted from frozen sperm samples with a QIAampDNA Mini kit (Qiagen). Exome sequences were efficiently enriched from 1 μg of genomic DNA using an Agilent Liquid Capture System (Agilent SureSelectHuman All Exon V6) according to the manufacturer’s protocol. The qualified genomic DNA was randomly fragmented to an average size of 180-280 bp by a Covaris S220 sonicator. DNA fragments were end-repaired, phosphorylated, and A-tail ligated at the 3’-end with the paired-end and single T base overhang adaptors (Illumina). After purification using AMPure SPRI beads from Agincourt, the size distribution and concentration of the resultant libraries were respectively determined using an Agilent 2100 Bioanalyzer and qualified using real-time PCR (2 nM). The DNA libraries were sequenced on an Illumina Hiseq 4000 using paired-end 150 bp reads.

### Statistics analysis

SPSS 19.0 software (SPSS Inc., USA) was used for statistical analysis, and groups was compared using two-tailed student’s t-tests. ANOVA was used to assess variation between groups.

## References

Arnott, D., Gawinowicz, M.A., Grant, R.A., Neubert, T.A., Packman, L.C., Speicher, K.D., Stone, K., and Turck, C.W. (2003). ABRF-PRG03: phosphorylation site determination. Journal of biomolecular techniques: JBT 14, 205–215.

Beltrao, P., Albanese, V., Kenner, L.R., Swaney, D.L., Burlingame, A., Villen, J., Lim, W.A., Fraser, J.S., Frydman, J., and Krogan, N.J. (2012). Systematic functional prioritization of protein posttranslational modifications. Cell 150, 413–425.

Bern, M., Cai, Y., and Goldberg, D. (2007). Lookup peaks: a hybrid of de novo sequencing and database search for protein identification by tandem mass spectrometry. Analytical chemistry 79, 1393–1400.

Bern, M., Kil, Y.J., and Becker, C. (2012). Byonic: advanced peptide and protein identification software. Current protocols in bioinformatics Chapter 13, Unit13 20.

Bhoj, V.G., and Chen, Z.J. (2009). Ubiquitylation in innate and adaptive immunity. Nature 458, 430–437.

Bregeon, D., and Doetsch, P.W. (2011). Transcriptional mutagenesis: causes and involvement in tumour development. Nature reviews Cancer 11, 218–227.

Chick, J.M., Kolippakkam, D., Nusinow, D.P., Zhai, B., Rad, R., Huttlin, E.L., and Gygi, S.P. (2015). A mass-tolerant database search identifies a large proportion of unassigned spectra in shotgun proteomics as modified peptides. Nature biotechnology 33, 743–749.

Creasy, D.M., and Cottrell, J.S. (2002). Error tolerant searching of uninterpreted tandem mass spectrometry data. Proteomics 2, 1426–1434.

Creasy, D.M., and Cottrell, J.S. (2004). Unimod: Protein modifications for mass spectrometry. Proteomics 4, 1534–1536.

Drezen, J.M., Gauthier, J., Josse, T., Bezier, A., Herniou, E., and Huguet, E. (2016). Foreign DNA acquisition by invertebrate genomes. Journal of invertebrate pathology.

Drummond, D.A., and Wilke, C.O. (2009). The evolutionary consequences of erroneous protein synthesis. Nature reviews Genetics 10, 715–724.

Fraley, C., and Raftery, A.E. (1998). How many clusters? Which clustering method? Answers via model-based cluster analysis. The Computer Journal 41, 578–588.

Fraley, C., Raftery, A.E., Murphy, T.B., and Scrucca, L. (2012). MCLUST Version 4 for R: Normal Mixture Modeling for Model-Based Clustering, Classification, and Density Estimation.

Garavelli, J.S. (2004). The RESID Database of Protein Modifications as a resource and annotation tool. Proteomics 4, 1527–1533.

Gasteiger, E., Gattiker, A., Hoogland, C., Ivanyi, I., Appel, R.D., and Bairoch, A. (2003). ExPASy: The proteomics server for in-depth protein knowledge and analysis. Nucleic acids research 31, 3784–3788.

Geng, Q., Ni, L., Ouyang, B., Hu, Y., Zhao, Y., and Guo, J. (2016). A Novel Testis-Specific Gene, Ccdc136, Is Required for Acrosome Formation and Fertilization in Mice. Reproductive sciences 23, 1387–1396.

Grunstein, M. (1997). Histone acetylation in chromatin structure and transcription. Nature 389, 349–352.

Hirsch, C., Gauss, R., Horn, S.C., Neuber, O., and Sommer, T. (2009). The ubiquitylation machinery of the endoplasmic reticulum. Nature 458, 453–460.

Hunter, T. (1995). Protein kinases and phosphatases: the yin and yang of protein phosphorylation and signaling. Cell 80, 225–236.

Kim, S., Gupta, N., Bandeira, N., and Pevzner, P.A. (2009). Spectral dictionaries: Integrating de novo peptide sequencing with database search of tandem mass spectra. Molecular & cellular proteomics: MCP 8, 53–69.

Liu, X., Wang, L., Zhao, K., Thompson, P.R., Hwang, Y., Marmorstein, R., and Cole, P.A. (2008). The structural basis of protein acetylation by the p300/CBP transcriptional coactivator. Nature 451, 846–850.

Lundgren, D.H., Martinez, H., Wright, M.E., and Han, D.K. (2009). Protein identification using Sorcerer 2 and SEQUEST. Current protocols in bioinformatics Chapter 13, Unit 13 13.

Mann, M., and Wilm, M. (1994). Error-tolerant identification of peptides in sequence databases by peptide sequence tags. Analytical chemistry 66, 4390–4399.

Mathias, R.A., Guise, A.J., and Cristea, I.M. (2015). Post-translational modifications regulate class IIa histone deacetylase (HDAC) function in health and disease. Molecular & cellular proteomics: MCP 14, 456–470.

Na, S., Bandeira, N., and Paek, E. (2012). Fast multi-blind modification search through tandem mass spectrometry. Molecular & cellular proteomics: MCP 11, M111 010199.

Na, S., and Paek, E. (2015). Software eyes for protein post-translational modifications. Mass spectrometry reviews 34, 133–147.

Perkins, D.N., Pappin, D.J., Creasy, D.M., and Cottrell, J.S. (1999). Probability-based protein identification by searching sequence databases using mass spectrometry data. Electrophoresis 20, 3551–3567.

Ribas de Pouplana, L. (2014). Not an inside job: non-coded amino acids compromise the genetic code. The EMBO journal 33, 1619–1620.

Sakamoto, T., Tanaka, T., Ito, Y., Rajesh, S., Iwamoto-Sugai, M., Kodera, Y., Tsuchida, N., Shibata, T., and Kohno, T. (1999). An NMR analysis of ubiquitin recognition by yeast ubiquitin hydrolase: evidence for novel substrate recognition by a cysteine protease. Biochemistry 38, 11634–11642.

Shen, P.S., Park, J., Qin, Y., Li, X., Parsawar, K., Larson, M.H., Cox, J., Cheng, Y., Lambowitz, A.M., Weissman, J.S., et al. (2015). Protein synthesis. Rqc2p and 60S ribosomal subunits mediate mRNA-independent elongation of nascent chains. Science 347, 75–78.

Smith, T.C., and Frank, E. (2016). Introducing Machine Learning Concepts with WEKA. Methods in molecular biology 1418, 353–378.

Tabb, D.L., Saraf, A., and Yates, J.R., 3rd (2003). GutenTag: high-throughput sequence tagging via an empirically derived fragmentation model. Analytical chemistry 75, 6415–6421.

Tak, Y.G., and Farnham, P.J. (2015). Making sense of GWAS: using epigenomics and genome engineering to understand the functional relevance of SNPs in non-coding regions of the human genome. Epigenetics & chromatin 8, 57.

Tsur, D., Tanner, S., Zandi, E., Bafna, V., and Pevzner, P.A. (2005). Identification of post-translational modifications by blind search of mass spectra. Nature biotechnology 23, 1562–1567.

Yang, J.H., Sklar, P., Axel, R., and Maniatis, T. (1995). Editing of glutamate receptor subunit B pre-mRNA in vitro by site-specific deamination of adenosine. Nature 374, 77–81.

Yates, J.R., 3rd (2015). Pivotal role of computers and software in mass spectrometry - SEQUEST and 20 years of tandem MS database searching. Journal of the American Society for Mass Spectrometry 26, 1804–1813.

Yi, C.H., Pan, H., Seebacher, J., Jang, I.H., Hyberts, S.G., Heffron, G.J., Vander Heiden, M.G., Yang, R., Li, F., Locasale, J.W., et al. (2011). Metabolic regulation of protein N-alpha-acetylation by Bcl-xL promotes cell survival. Cell 146, 607–620.

